# Immediate effect of quadri-pulse stimulation on human brain microstructures

**DOI:** 10.1101/2023.04.20.537631

**Authors:** Ikko Kimura, Masamichi J Hayashi, Kaoru Amano

**Affiliations:** Center for Information and Neural Networks (CiNet), Advanced ICT Research Institute, National Institute of Information and Communications Technology, Suita 565-0871, Japan; Graduate School of Frontier Biosciences, Osaka University, Suita 565-0871, Japan; Graduate School of Information Science and Technology, The University of Tokyo, Tokyo 113-8656, Japan

**Keywords:** diffusion-tensor imaging, quadri-pulse stimulation, repetitive transcranial magnetic stimulation, fractional anisotropy, mean diffusivity

## Abstract

Several studies have implied that human brain microstructures can change immediately after a behavioral training. However, since widespread regions are involved in behavioral training, it remains unclear whether the microstructure in the living human brain changes immediately after the change in activity of a specific brain area. Hence, we aimed to examine whether the microstructures in the human brain change after the increase and decrease in the specific brain activity by repetitive transcranial magnetic stimulation, namely quadri-pulse stimulation (QPS). Right-handed healthy adults underwent both the excitatory (QPS5) and inhibitory (QPS50) QPS protocols over the left M1. Before and after QPS, diffusion MRI and resting-state fMRI scans were collected to detect any microstructural (fractional anisotropy [FA] and mean diffusivity [MD] values) and functional (functional connectivity between the bilateral M1) changes after QPS5 and QPS50. As a result, we observed no statistically significant change in FA or MD values after either QPS5 or QPS50 in cerebral cortex. This suggests that the brain activity change in widespread area is required to induce microstructural change immediately.

## Introduction

Neural activity shapes the underlying microstructure of its activity. Several animal studies have suggested that repeated activation of neurons increases dendritic spines^1^ or enlargens astrocytes^2^ in the gray matter, and induces myelination in the white matter near the activated neurons^3^ (see Sampaio-Baptista and Johansen-Berg (2017)^4^ for a review). Several human studies have also observed microstructural changes in the brain a day after neural intervention^5–7^, suggesting that microstructural plasticity may also occur in the adult human brain. Since brain functions can change in both normal (e.g., behavioral training) and abnormal (e.g., stroke, brain tumor, or epilepsy) situations, it is crucial to elucidate how functional changes are related to microstructural changes.

While most previous studies have examined microstructural plasticity more than a day after neural interventions^4^, several studies suggest that microstructural changes can also occur immediately after the intervention^8^. Sagi et al. showed that the microstructural property revealed by diffusion MRI (dMRI) were modulated immediately after a single session of a spatial memory task^2^. A recent study also showed that microstructural changes can be observed immediately after 45 min of motor training^9^. Although this type of microstructural plasticity in the ultra-acute phase is also important in unraveling its association with functional changes, behavioral training changes the brain activity in widespread areas^10^, rather than in a specific area. Therefore, it is still unclear whether the microstructure changes immediately after the chage in brain activity of a specific region. The comparison of the effect induced by a facilitation and suppression of a brain activity is also crucial because not only facilitation (e.g., epilepsy) but also suppression (e.g., stroke or brain tumor) of the brain activity in a specific region can occur in clinical situations.

Brain activity can be bidirectionally modulated by repetitive transcranial magnetic stimulation (rTMS). This method can noninvasively facilitate or suppress the brain activity of a specific region to examine the direct effects of neural intervention^11^. A few previous studies have investigated microstructural changes immediately after either the excitatory^14^ or inhibitory^15–17^ protocol of conventional rTMS with inconsistent results. Instead of conventional rTMS, we here used quadri-pulse stimulation (QPS), which is one of rTMS protocols, and induces strong and robust after-ffect^12, 13^. We tested whether the microstructural properties change immediately after both QPS5 and QPS50, and that the degree of change might differ between after QPS5 and QPS50. This knowledge is important not only for clarifying the susceptibility of structural plasticity in the ultra-acute phase but also for elucidating the potential of rTMS as a method to modulate the microstructural properties of the human brain.

## Results

The three-day-long experiments were performed to evaluate the changes in both microstructural and functional properties in the human brain after QPS (Figure 1). On Day 1, task functional MRI (fMRI) during a finger tapping task and structural MRI were obtained for identifying the primary motor cortex (M1) responsible for the right index finger and for the spatial normalization in group analysis, respectively. On Day 2 or 3, QPS5 or QPS50 over the left M1 was applied, and before and after the QPS session, dMRI and resting-state fMRI (rsfMRI) were collected to detect the microstructural and functional changes, respectively.

**Figure 1.**
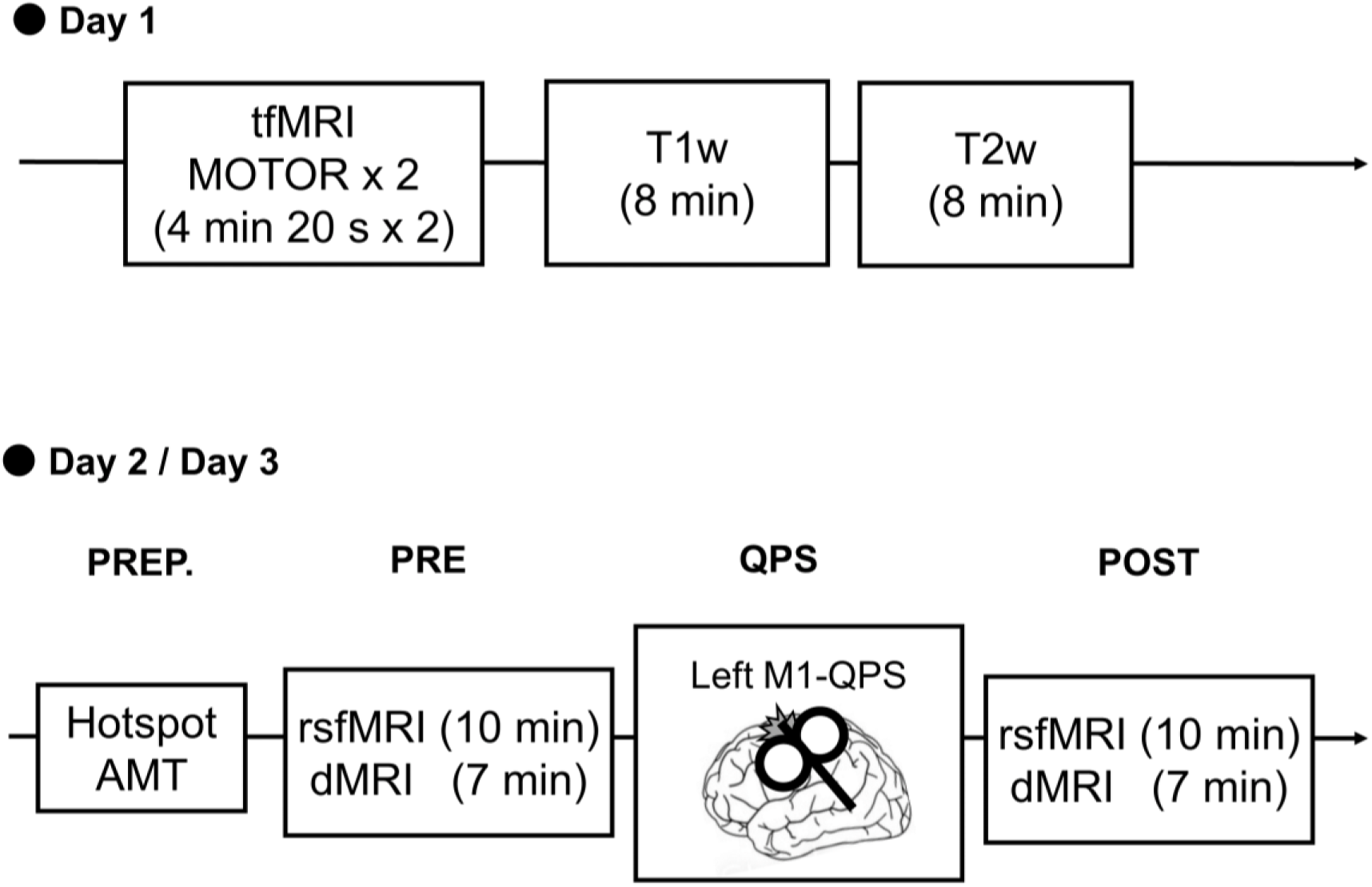
Overview of the experimental protocol to assess the microstructural changes after QPS over the primary motor cortex (M1). On Day 1, task fMRI (tfMRI) data were acquired during a finger tapping task (MOTOR) to localize the left M1, while T1w and T2w images were obtained to spatially normalize the diffusion MRI (dMRI) and resting-state fMRI (rsfMRI) data. On Days 2 and 3, either QPS5 or QPS50 was performed. First, stimulus location (hotspot for left M1) and stimulus intensity (active motor threshold; AMT) were defined in the preparation step (PREP.). Before and after QPS, participants underwent resting-state fMRI (rsfMRI) and diffusion MRI (dMRI) scans.

All 16 healthy adults completed the three-day-long experiment without any adverse events. One session in QPS5 failed to be recorded by a neuronavigation system, and two sessions of task fMRI could not be performed because of the difficulty in presenting the visual stimuli for the timing of the finger tapping. Paired t-tests revealed that there were no significant differences in active motor threshold (AMT; *t* = 0.40, *P* = 0.70; Figure 2A) or stimulus intensity for QPS (*t* = 0.48, *P* = 0.64; Figure 2B) between the QPS5 and QPS50 conditions. There were no significant differences in the quality of the QPS-session (mean distance error, *t* = 0.77, *P* = 0.46, Figure 2C; mean angle error, *t* = −1.37, *P* = 0.19, Figure 2D; mean twist error, *t* = −0.56, *P* = 0.59, Figure 2E).

**Figure 2.**
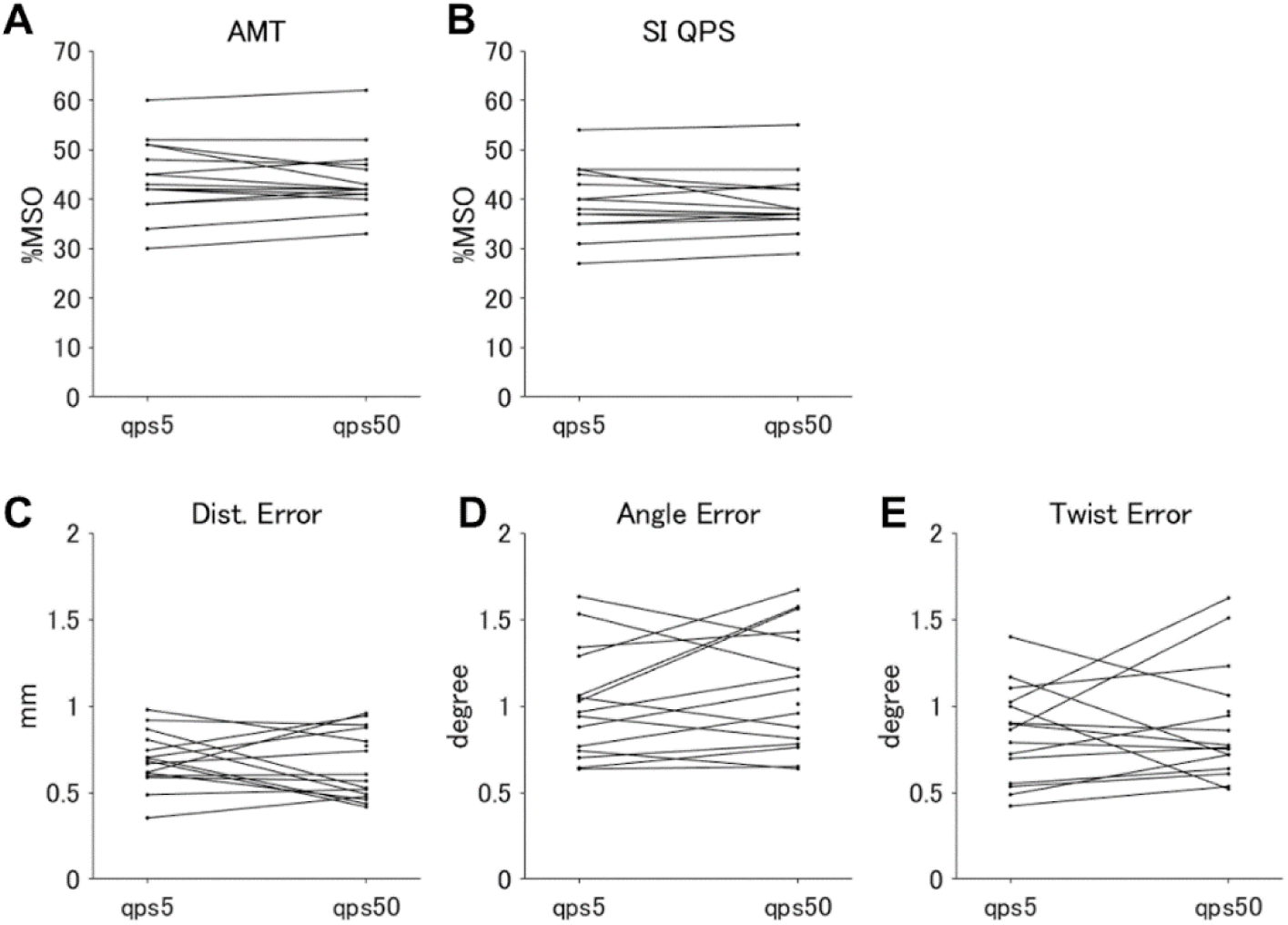
(A–B) Comparison of baseline neurophysiological features between the QPS5 and QPS50 conditions. The active motor threshold (AMT) and stimulus intensity (SI) for applying QPS were shown in (A) and (B), respectively. The vertical axes in the upper panel illustrate the percentage of maximum stimulator output (% MSO). (C–E) Comparison of the quality in applying QPS between the QPS5 and QPS50 conditions. The mean distance error (mm; C), the mean angle error (degree; D), and the mean twist error (degree; E) in each session was shown.

### The change of the microstructural property after quadri-pulse stimulation

The changes in the microstructural properties of the gray matter in both cerebral cortex and subcortical regions after QPS5 or QPS50 were evaluated by comparing the metrics derived from dMRI (fractional anisotropy [FA] and mean diffusivity [MD] values) before and after QPS5 or QPS50. In cerebral cortex, we found no significant changes after both QPS5 and QPS50 in both FA (QPS5: Bayes factor, Figure 3A, Effect size, Supplementary Figure 1A; QPS50: Bayes factor, Figure 3B, Effect size, Supplementary Figure 1B) and MD (QPS5: Bayes factor, Figure 4A, Effect size, Supplementary Figure 2A; QPS50: Bayes factor, Figure 4B, Effect size, Supplementary Figure 2B) values. In contrast, we found significant decrease in MD value in the bilateral cerebellum after both QPS5 (Figure 5A) and QPS50 (Figure 5B). Except for this change, no significant change was detected in subcortical regions in both FA (QPS5: Bayes factor, Figure 3D, Effect size, Supplementary Figure 1D; QPS50: Bayes factor, Figure 3E, Effect size, Supplementary Figure 1E) and MD (QPS5: Bayes factor, Figure 4D, Effect size, Supplementary Figure 2D; QPS50: Bayes factor, Figure 4E, Effect size, Supplementary Figure 2E) values. There was no significant difference in the changes in FA (Bayes factor, Figures 3C and F; Effect size, Supplementary Figures 1C and F) and MD (Bayes factor, Figures 4C and F; Effect size, Supplementary Figures 2C and F) values between the QPS5 and QPS50 conditions.

**Figure 3.**
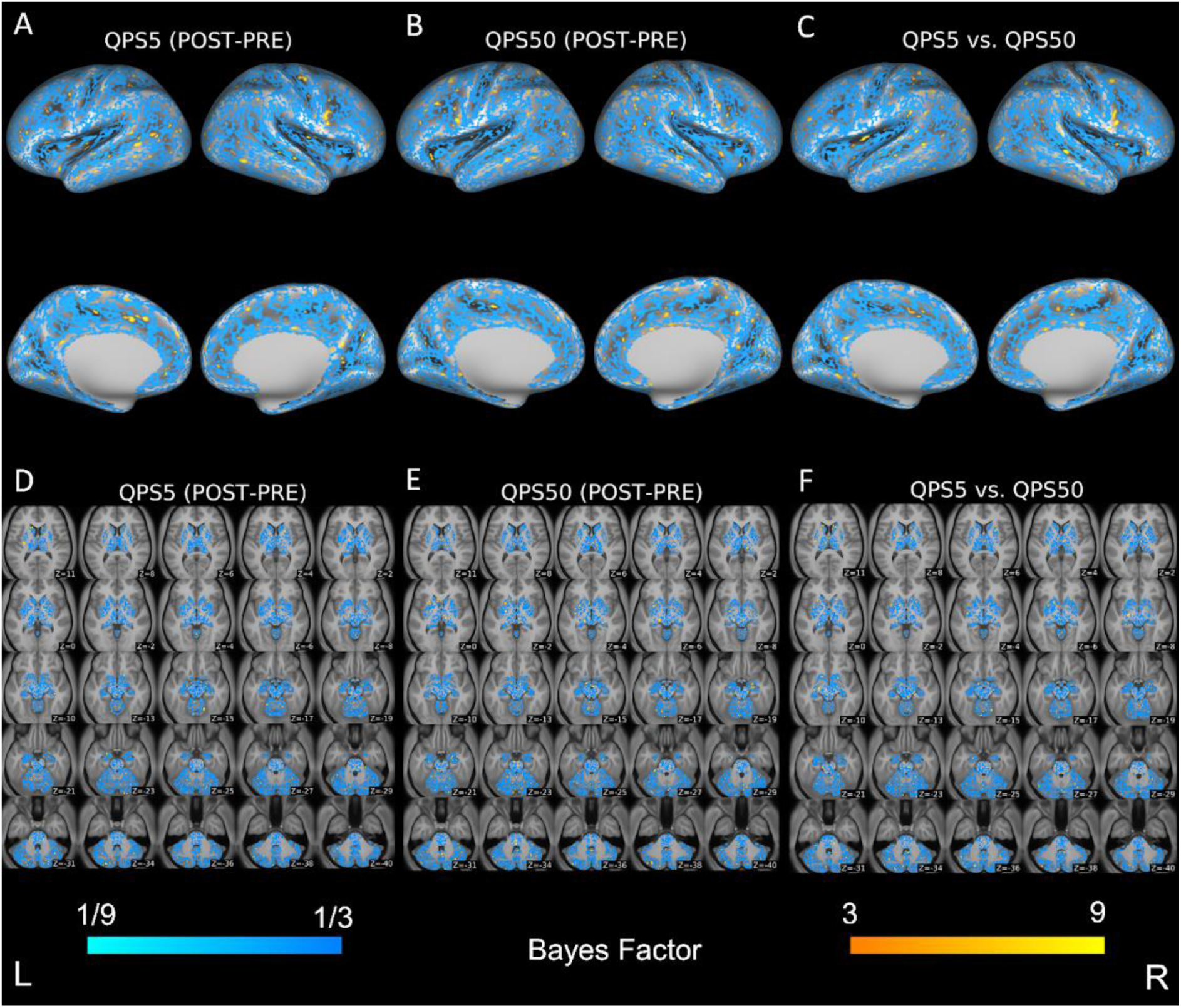
Differences in FA values of the cerebral cortex and subcortical regions between pre- and post-QPS in QPS5 (A and D) and QPS50 (B and E) conditions, and those in changes of FA values across conditions (C and F). The upper panel (A–C) shows the result of surface-based analysis on the cerebral cortex, while the lower panel (D–F) shows that of voxel-based analysis in the subcortical regions. Areas in blue illustrate that the Bayes factor was smaller than ⅓ in the comparison of the FA values before and after QPS or the change of the FA values across conditions (i.e., supports the hypothesis that the difference was zero). In contrast, those in yellow show those values are higher than 3. Axial slices in (D–F) are shown in accordance with neurological conventions (the left side of the image is of the left of the brain) and are displayed in the MNI coordinates from z = 11 (top left) to z = −40 (bottom right).

**Figure 4.**
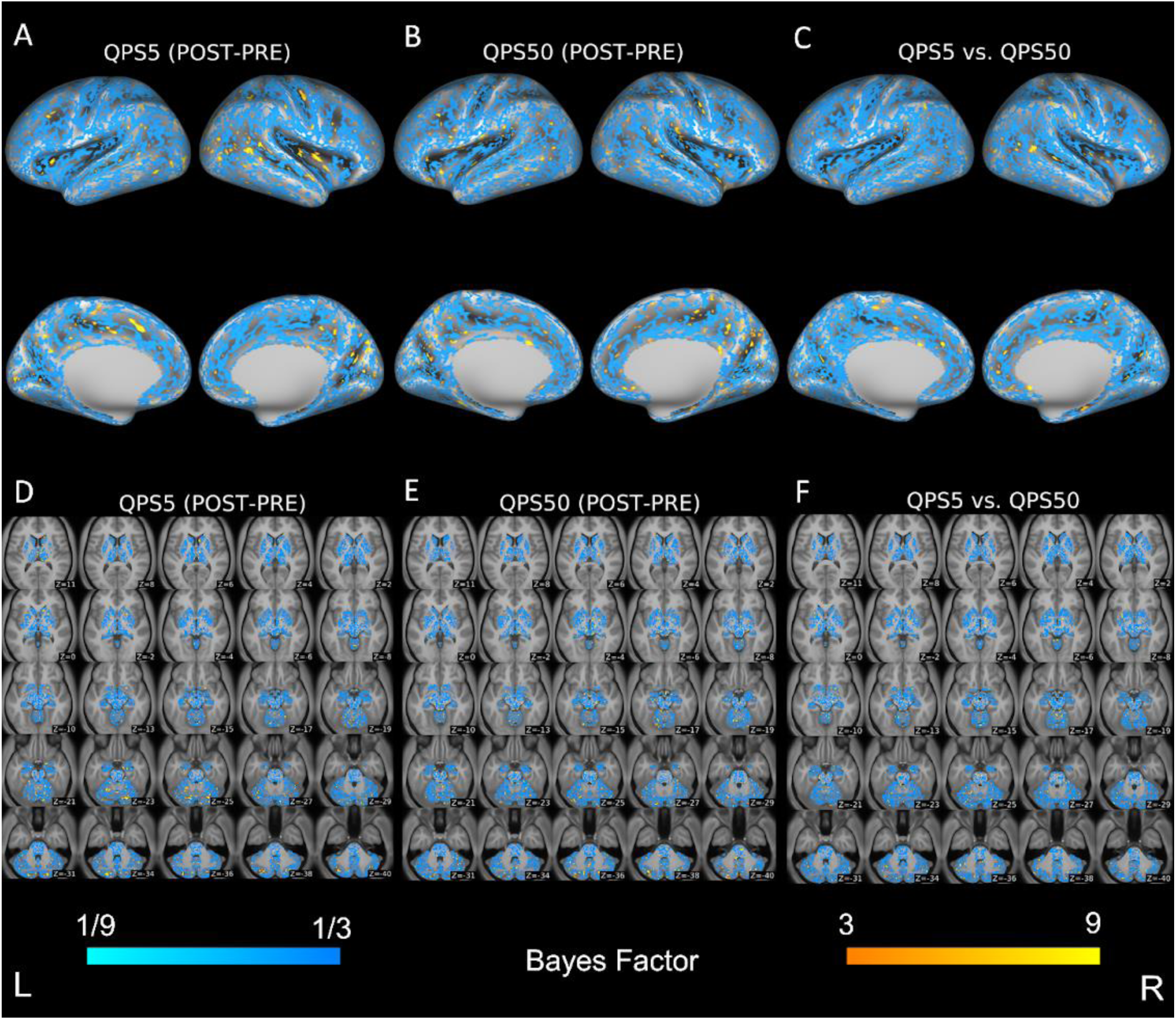
Differences in MD values of the cerebral cortex and subcortical regions between pre- and post-QPS in QPS5 (A and D) and QPS50 (B and E) conditions, and those in changes of MD values across conditions (C and F). The upper panel (A–C) shows the result of surface-based analysis on the cerebral cortex, while the lower panel (D–F) shows that of voxel-based analysis in the subcortical regions. Areas in blue illustrate that the Bayes factor was smaller than ⅓ in the comparison of the MD values before and after QPS or the change of the MD values across conditions (i.e., supports the hypothesis that the difference was zero). In contrast, those in yellow show those values are higher than 3. Axial slices in (D–F) are shown in accordance with neurological conventions (the left side of the image is of the left of the brain) and are displayed in the MNI coordinates from z = 11 (top left) to z = −40 (bottom right).

**Figure 5.**
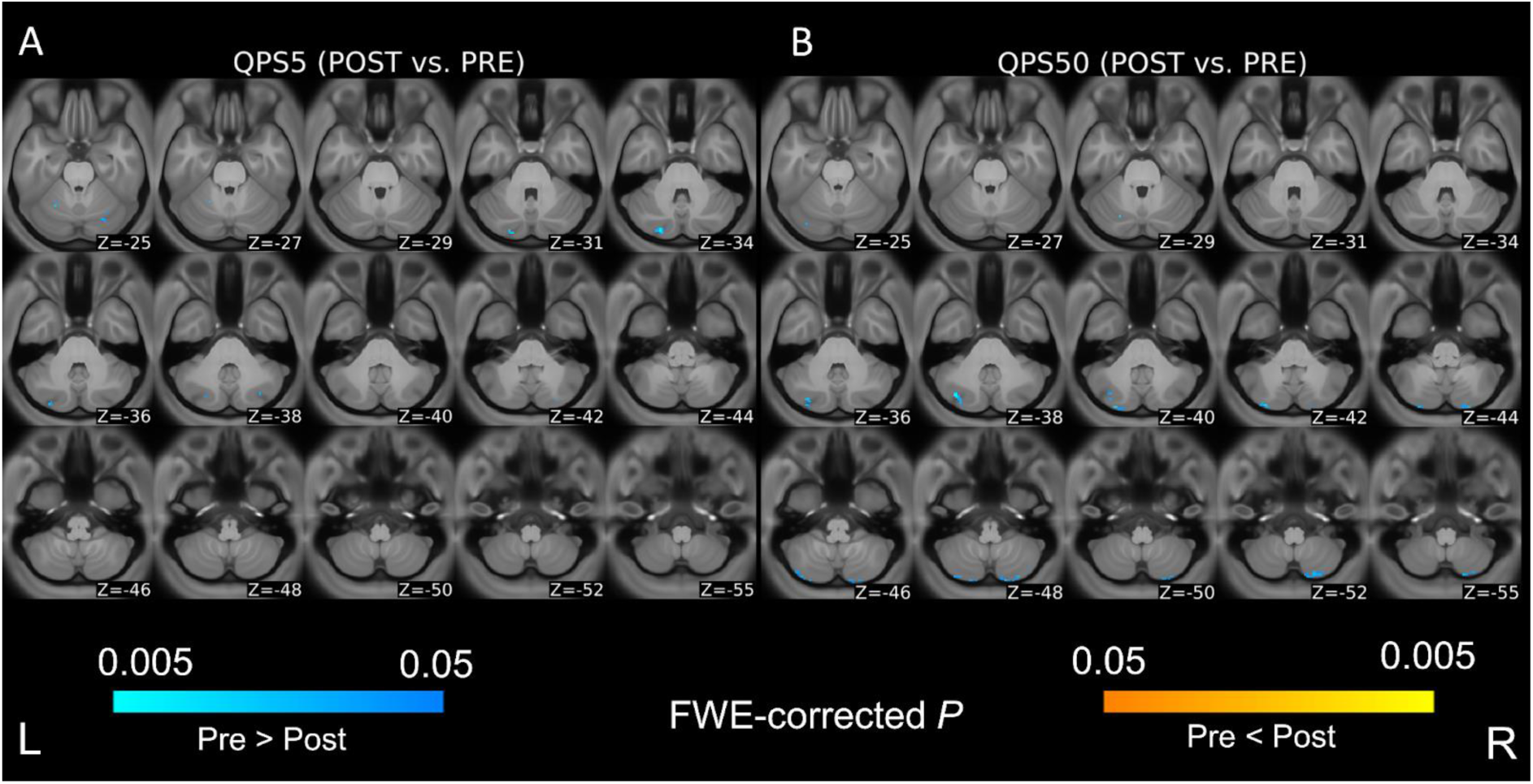
Differences in MD values between pre- and post-QPS in QPS5 (left panel) and QPS50 (right panel) conditions. Areas in blue show that MD was significantly decreased after QPS, while areas in yellow indicate that the value was significantly increased after QPS. Axial slices are shown in accordance with neurological conventions (the left side of the image is of the left of the brain) and are displayed in the MNI coordinates from z = −25 (top left) to z = −55 (bottom right).

We then assessed the changes in the microstructural properties of the white matter after QPS5 and QPS50. Tract-based spatial statistics (TBSS) revealed no significant changes after QPS5 or QPS50 in both FA (QPS5: Bayes factor, Supplemetary Figure 3A, Effect size, Supplementary Figure 4A; QPS50: Bayes factor, Supplemetary Figure 3B, Effect size, Supplementary Figure 4B) and MD (QPS5: Bayes factor, Supplemetary Figure 5A, Effect size, Supplementary Figure 6A; QPS50: Bayes factor, Supplemetary Figure 5B, Effect size, Supplementary Figure 6B) values of the white matter. No significant differences in the changes in the FA (Bayes factor, Supplemetary Figure 3C; Effect size, Supplementary Figure 4C) and MD (Bayes factor, Supplemetary Figure 5C; Effect size, Supplementary Figure 6C) values were observed in the white matter between the QPS5 and QPS50 conditions.

### Relationship between changes in functional connectivity in the stimulated region and the microstructural properties of the brain after quadri-pulse stimulation

Although we found the no significant microstructural changes after QPS in most regions, to investigate whether inter-individual variability in microstructural changes in the brain after QPS is related to functional changes, we first compared the functional connectivity (FC) of the stimultad region (left M1) assessed with seed-based correlation analysis of rsfMRI before and after QPS5 or QPS50. After QPS5, the FC of the left M1 was significantly decreased in the right dorsomedial prefrontal cortex, ventral premotor cortex, precentral gyrus, and postcentral gyrus (Supplementary Figure 7A), whereas it was significantly increased in the bilateral cerebellum (Supplementary Figure 7C). In contrast, after QPS50, there was a significant decrease in the FC of the left M1 in the left supplementary motor area, left precentral gyrus, and bilateral postcentral gyrus (Supplementary Figure 7B), whereas no significant alteration was observed in the FC between the left M1 and the subcortical structures (Supplementary Figure 7D).

Since the previos study showed that the change in FC between the left and right M1 was negatively correlated with the degree of the after-effect of QPS over the left M1^18^, we then investigated the change in the FC between these two regions with region of interest (ROI) to ROI analysis. The FC between the left and right M1 decreased after QPS5 (*t* = −2.60, *P* = 0.020), whereas the FC was not significantly modulated after QPS50 (*t* = −1.23, *P* = 0.24) (Figure 6). There were no significant differences between the QPS5 and QPS50 conditions regarding the change in FC between these two regions after QPS (*t* = −0.80, *P* = 0.44).

**Figure 6.**
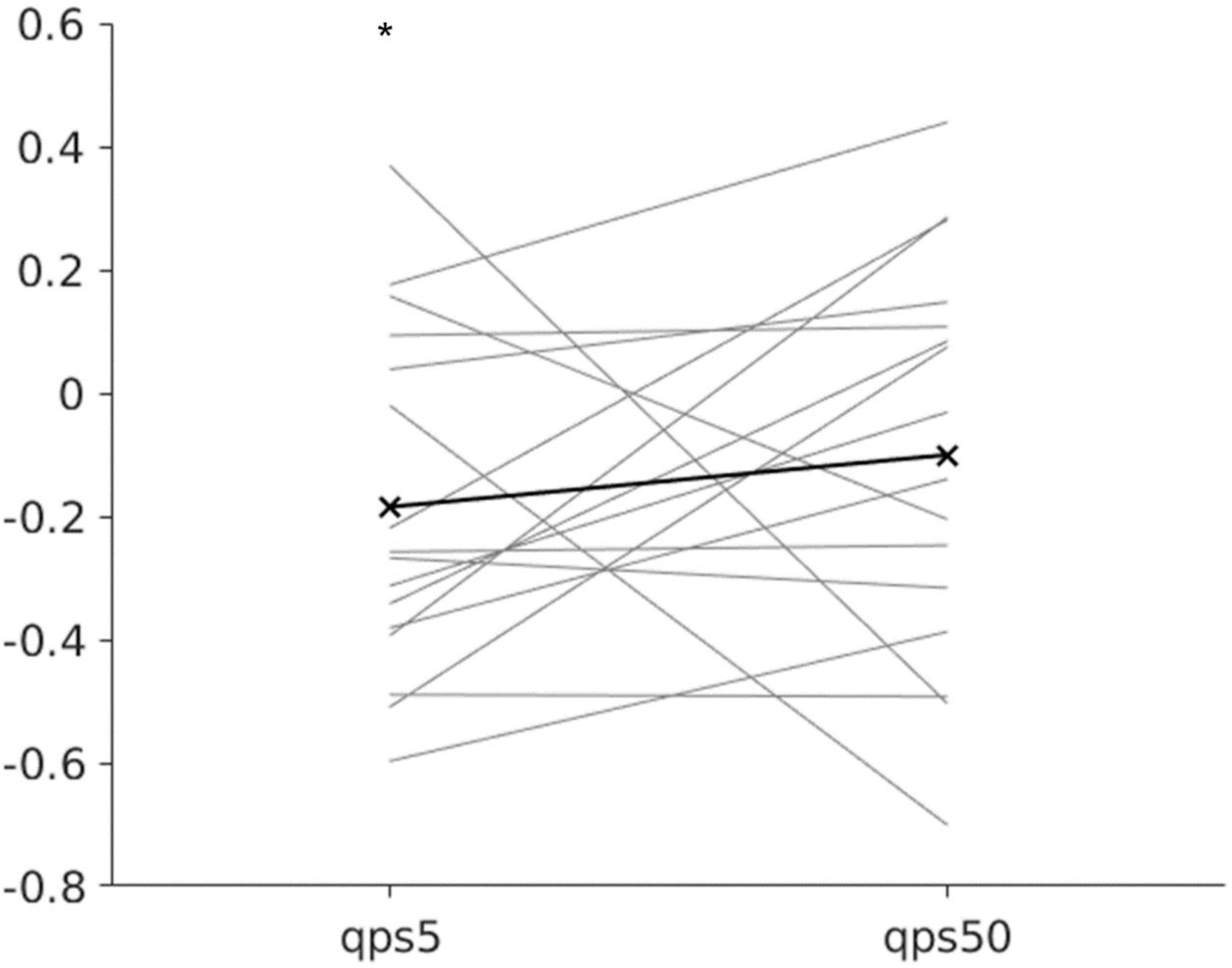
Changes in the functional connectivity (FC) between the left and right M1 after QPS. Thick black line bordered by crosses indicates the mean FC change across participants. The figure on the left denotes the change after QPS5, while that on the right denotes that after QPS50. The y-axis indicates the change in the FC after QPS. * shows the change was statistically significant (*P* < 0.05).

The correlation between the change in FA or MD values and the change in FC between the left and right M1 after QPS was investigated. Regression analysis revealed that there were no significant correlations between the change in FA or MD value and the change in FC between the left and right M1 after either QPS5 or QPS50. To confirm that the size of the ROI did not change the results, Pearson’s correlation coefficient between the change in FC defined by a five mm radius (original) and that defined by a two mm radius or that defined by an eight mm radius in both QPS5 and QPS50 conditions was calculated. As a result, we found that all of the changes were significantly and positively correlated with each other, suggesting that the size of the ROI did not affect the change in FC between the left and right M1 (QPS5 [five mm radius vs. eight mm radius], rho = 0.94, *P* < 0.001; QPS5 [five mm radius vs. two mm radius], rho = 0.97, *P* < 0.001; QPS50 [five mm radius vs. eight mm radius], rho = 0.94, *P* < 0.001; QPS50 [five mm radius vs. two mm radius], rho = 0.92, *P* < 0.001). Additionally, a significant correlation was not observed between the mean decrease in the MD value and the mean change in FC with the left M1 in the clusters of the bilateral cerebellum, whose MD values significantly decreased after QPS5 (rho = −0.21, *P* = 0.43) or QPS50 (rho = −0.35, *P* = 0.18).

### Possible confounding factors both within and across the session

To detect the confounding factors affecting the change after QPS5 or QPS50 or the difference in the change between the QPS5 and QPS50 conditions, we compared the quantitative measures of image quality between pre- and post-QPS and the change in these measurements between the QPS5 and QPS50 conditions. Regarding rsfMRI, Wilcoxon signed-rank tests revealed no significant changes in the Stanford Sleepiness Scale score (sleepiness) after QPS5 or QPS50 (QPS5, *W* = 65.50, *P* = 0.41; QPS50, *W* = 63.00, *P* = 0.61; Figure 7A). Though no significant change in the mean framewise displacement (FD), which is an index for the degree of a participant’s movement during a scan, was found after QPS5 (*t* = 1.24, *P* = 0.23, Figure 7B), there was a significant decrease after QPS50 (*t* = 2.69, *P* = 0.017, Figure 7B). No significant changes were found in the image quality of dMRI after either QPS5 (mean absolute motion, *t* = −0.99, *P* = 0.34, Figure 7C; mean relative motion, *t* = −0.52, *P* = 0.61, Figure 7D) or QPS50 (mean absolute motion, *t* = 0.51, *P* = 0.62, Figure 7C; mean relative motion, *t* = 0.78, *P* = 0.45, Figure 7D). There was no significant difference between the QPS5 and QP50 conditions in the change in all these measurements within the session (Stanford Sleepiness Score, *W* = 49.50, *P* = 0.80, Figure 7A; mean FD, *t* = 0.99, *P* = 0.34, Figure 7B; mean absolute motion in dMRI, *t* = 1.23, *P* = 0.24, Figure 7C; mean relative motion in dMRI, *t* = 0.83, *P* = 0.42, Figure 7D).

**Figure 7.**
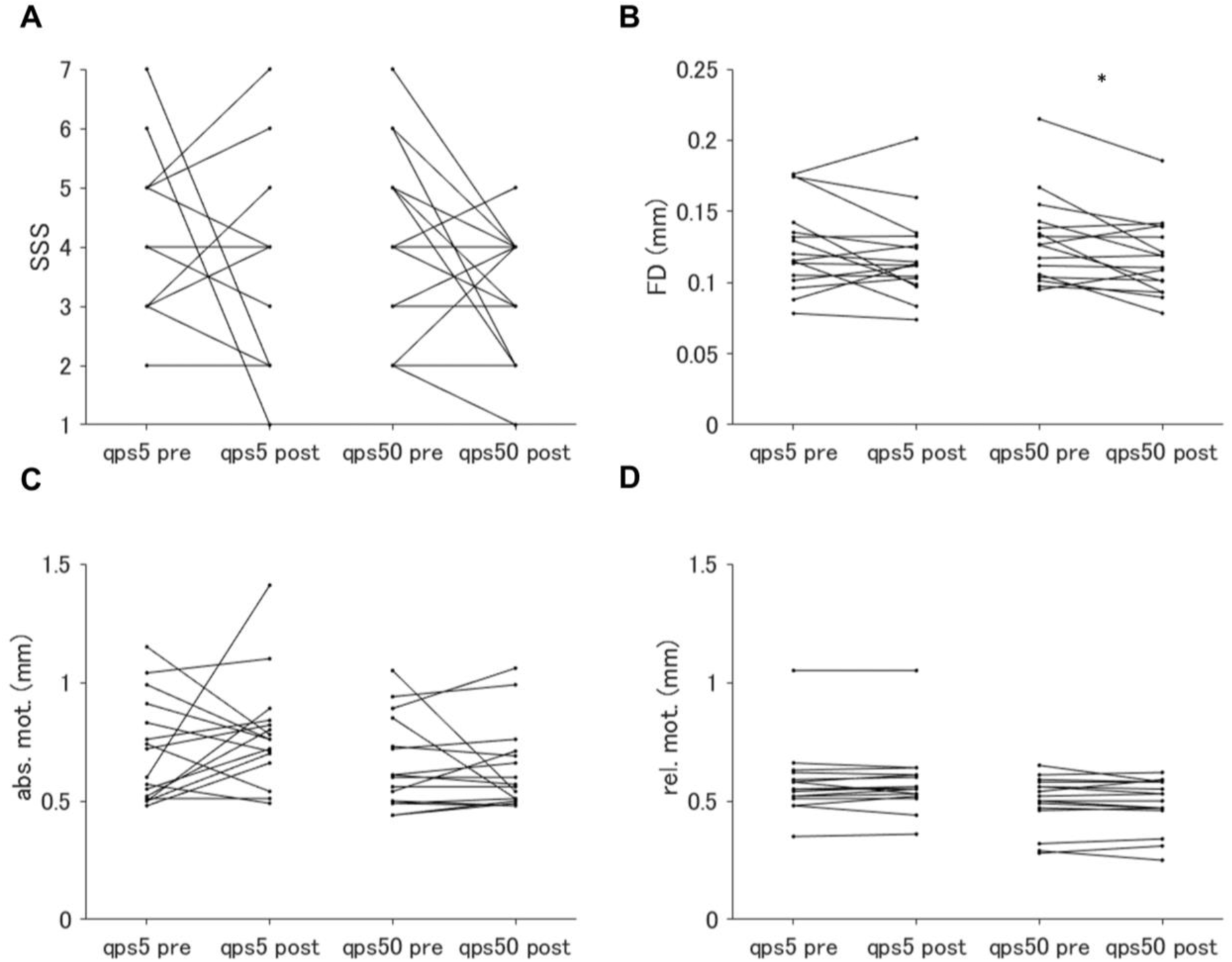
Comparisons of the quantitative measurements on the quality of resting-state fMRI (A–B) and diffusion MRI (dMRI; C–D) measured pre- and post-QPS in QPS5 and QPS50 conditions. (A) and (B) show Stanford Sleepiness Scale (SSS) and mean framewise displacement (FD; mm), respectively, while (C) and (D) illustrate mean absolute and relative motion (mm) during dMRI scan. * indicates that the comparison was significantly different according to the paired t-test.

Since the mean FD can be a confounding factor when comparing the FC of the left M1 before and after QPS, we also analyzed the FC of the left M1, regressing out the effect of mean FD. As a result, we confirmed that the mean FD did not severely affect the results related to the change in the FC of the left M1 (Supplementary Results).

## Discussion

Here, we studied whether QPS5 or QPS50 over the left M1 changes the microstructural properties of the cerebral cortex, subcortical regions, and white matter of the human brain. As a result, no significant changes were found in the FA or MD values after either QPS5 or QPS50 except for the significant decrease in the MD value in the bilateral cerebellum after both QPS5 and QPS50. No significant correlations were observed between the change in microstructural properties and that in FC between the left and right M1 (the functional change of QPS) in after either QPS5 or QPS50.

In cerebral cortex, no significant change was detected in the FA or MD values after either QPS5 or QPS50 and no significant correlation was observed between the change in these values and the functional changes in QPS. Given that FA and MD values are sensitive biomarkers for changes in the microstructural properties of both neurons^19, 20^ and glial cells^2, 21, 22^ in the brain, the current results suggest that the microstructural properties in the cerebral cortex were not modulated immediately after both the excitatory and inhibitory protocols of QPS. Our results are consistent with previous studies reporting that the microstructural properties were not significantly changed after either the excitatory^14^ or inhibitory^16^ protocol of conventional rTMS. Given that the microstructural property changed immediately after a single session of behavioral training^2, 9^, we speculate that widespread brain activity change rather than the change in brain activity of a specific brain region evokes the microstural change immediately after the neural intervention in the living human brain.

In contrast, two previous studies reported microstructural alterations in the human brain immediately after rTMS. While one study observed that MD values were significantly decreased in the stimulated region^17^, another study reported that these values were significantly increased in both the stimulated and unstimulated regions^15^ after 1 Hz conventional rTMS, which decreases the brain activity of the stimulated region. These differences might be partially because of the differences in the rTMS protocols. In previous studies, the stimulus intensity was 90% resting motor threshold (RMT)^15, 17^, which was stronger than that used in the current study (90% AMT). Moreover, the neural mechanisms evoked by QPS and 1 Hz rTMS are different. While short-interval intracortical inhibition, which measures the function of inhibitory interneurons^23^, is decreased by 1 Hz rTMS^24^, it is not modulated by QPS^12^. Animal studies confirmed that the number of calbindin-expressing inhibitory interneurons decreased after 1 Hz rTMS^25^, implying that the change in the microstructural properties induced by the 1 Hz rTMS might reflect the microstructural changes in inhibitory interneurons. Therefore, the lack of significant microstructural alterations in the cerebral cortex in the current study may be specific to QPS.

The MD values of the bilateral cerebellum decreased after both QPS5 and QPS50, and the change in the MD value was not significantly different across the conditions. The cerebellum is important for shaping the internal model of action for precise movement^26^. Moreover, one study reported that patients with cerebellar degeneration failed to perform a reaching task in a new environment, suggesting that the cerebellum also plays a critical role in adapting to new environments during movement^27^. Together with the results of the current study, these results might imply that the microstructural properties of the bilateral cerebellum were changed to adapt to the change in the brain activity of the left M1. However, these results must be interpreted with caution. This is because the changes in these properties were similar between QPS5 and QPS50 conditions and were not correlated with the functional change, implying that the change in the microstructural properties in this study might be unspecific to the after-effect of QPS. Also, voxel-based analysis with dMRI, as opposed to the surface-based analsysis used in cerebral cortex, has been reported to be susceptible to registration errors^28^. Future animal studies or MRI studies with a high spatial resolution are required to verify whether these changes actually occur.

The FA or MD values of white matter did not change after either QPS5 or QPS50. This result is consistent with a previous study showing that white matter volume was not significantly modulated immediately after continuous theta-burst stimulation, which is an inhibitory patterned rTMS protocol, to the temporal area^29^. In contrast, several studies have reported the chnage in the microstructural properties of white matter 24 h after neuromodulation interventions, such as neurofeedback training^6^ and cortico-cortico paired associative stimulation^5^. These differences might be due to the differences in time evaluated after the intervention because several animal studies have shown that oligodendrogenesis or myelination can be observed more than one hour after stimulation^4^. Since one study reported that oligodendrogenesis can occur several hours after repetitive neural activation^3^ and the after-effect of QPS lasts for more than several hours^12^, future studies are required to investigate the microstructural changes several hours after QPS.

In the current study, the FC between the left and right M1 decreased after QPS5 but was not significantly modulated after QPS50. These results partially contradict the previous study showing that the FC between these two regions decreased after QPS5, whereas it increased after QPS50^18^. These differences could be explained by the time of measurement after QPS: immediately after QPS in the current study and 30 minutes after QPS in the study by Watanabe et al.^18^. Given that bidirectional changes in motor evoked potential (MEP) amplitude^13^ after QPS over the left M1 (increase after QPS5 and decrease after QPS50) were clearly observed 30 min after QPS^13^, the changes in FC between the left and right M1 might also be observed more clearly 30 min after QPS than immediately after QPS. Also, while the after-effect of QPS was assessed with the change in FC between the left and right M1^32^, mainly due to time constraints, MEP will more accurately reveal the association between microstructural and functional changes after QPS.

One of the future directions of this study would be to assess the microstructural changes with different modalities. In the current study, the microstructural properties were assessed using the dMRI-derived metrics. Although these metrics are useful in exploring changes in the microstructural properties of the human brain^2, 19, 21, 30^, a previous study detected microstructural alterations after neuromodulation interventions using magnetization transfer imaging^5^, which is a sensitive biomarker of myelin. Another study implied that the microstructural metrics derived from differe modalities might reflect different types of microstructural characteristics^31^. Therefore, microstructural indicators obtained with neuroimaging modalities other than dMRI might detect the change in microstructural property after QPS.

Another interesting future direction is to assess the change in astrocytes or microglial cells induced by rTMS. Several animal studies have suggested that the activity of glial cells is modulated by rTMS^33^. Because these functional properties can also be noninvasively evaluated in the human brain using PET^34^, it would be interesting to explore the correlations between functional changes in glial cells and microstructural changes in the human brain after QPS.

## Conclusion

There were no significant microstructural changes in the human brain immediately after QPS5 or QPS50 except for the decrease in MD values in the bilateral cerebellum. These results suggest that the change in widespread brain activity is required for the microstructure in the human brain to change immediately after the intervention. Future studies are needed to examine microstructural changes at different time points after the intervention and to assess these changes using different modalities.

## Methods

The study consisted of three days of experiments (Figure 1). On Day 1, task fMRI data were collected to localize the M1 responsible for the right first dorsal interosseous (FDI) muscle. Structural MRI data were collected on the same day (Figure 1A). On Days 2 and 3, the participants underwent either QPS5 or QPS50. Before and after QPS, rsfMRI and dMRI data were obtained to assess the functional and microstructural changes in QPS, respectively (Figure 1B). Days 2 and 3 were both performed in the afternoon, from 1 pm to 3 pm, and were separated for more than a week to avoid any cumulative effects. QPS5 and QPS50 were randomly assigned to Days 2 and 3 and the order was counterbalanced across participants.

### Participants

Sixteen healthy adult volunteers (eight males; age range 20–25 years, mean 22.7 years, standard deviation (SD) 1.4 years) participated in this study. All participants were right-handed according to the Edinburgh handedness inventory (mean score 95.05, SD 9.8) and reported no history of neurological or psychiatric disorders. The experiments were approved by the Institutional Ethics and Safety Committees of the National Institute of Information and Communications Technology, and were performed in accordance with the Declaration of Helsinki. Informed consent was obtained from all participants after providing them with a full explanation of the study protocols and objectives.

### Quadri-pulse Stimulation

Participants underwent QPS5 or QPS50 over the left M1, targeting the right FDI muscle, using the same procedure as described in a previous study^13^. In brief, the motor hotspot for the right FDI muscle was first determined, and from this hotspot, the AMT was defined (evoking MEPs larger than 100 mV in 5 out of 10 trials with mild contraction of the right FDI). The QPS was then applied with a burst of four monophasic pulses every 5 s for 30 min, and the stimulus intensity was set at 90% of the AMT. The inter-pulse interval for QPS5 was 5 ms, while that for QPS50 was 50 ms. These procedures were performed with a DuoMAG MP-Quad (Deymed Diagnostic s.r.o., Hronov, Czech Republic) equipped with a butterfly-shaped 70-mm air-cooling coil (DuoMAG 70BF Air Cooled Coil; Deymed Diagnostic s.r.o., Hronov, Czech Republic), and MEPs were collected using a Brainsight surface-electromyogram (Rogue Research Inc., Montreal, Canada). To quantitatively assess the stability of the coil position during QPS, the position was recorded using a Brainsight (Rogue Research Inc., Montreal, Canada). The mean distance error of the coil position and the mean angle or twist error of the coil handle relative to the target position of the coil were calculated in each QPS session to evaluate the quality of applying QPS in each session.

### Image Acquisition

On Day 1, two sessions of task fMRI data during a finger tapping task and structural MRI data were acquired using a Siemens PrismaFit 3T scanner equipped with a 32-channel array head coil (Siemens, Erlangen, Germany). On Days 2 and 3, rsfMRI and dMRI data before and after QPS were obtained using a Siemens Vida 3T scanner equipped with a 64-channel array head-neck coil (Siemens, Erlangen, Germany).

To identify the left M1 responsible for the right FDI and define its location as the seed region for the subsequent seed-based correlation analysis, two sessions of fMRI data were collected while participants tapped their right or left index fingers, as described in a previous study^18^. Each session consisted of six blocks of 20 s with an inter-block interval of 20 s and took 4 min 20 s in total. In each block, participants were required to tap their left or right index fingers synchronized with the center red circle (radius 1 °) flashing at 1 Hz. These fMRI data were collected with a gradient-echo echo-planar sequence, and the slices were collected in an interleaved order. The other acquisition parameters were as follows: flip angle = 60 °; voxel size = 2 × 2 × 2 mm; matrix size = 108 × 108 × 78; Multiband Acceleration Factor = 6, phase-encoding direction = anterior-posterior; TR = 1000 ms; and TE = 30 ms. A pair of B0 field maps in both phase-encoding directions (i.e., anterior–posterior and posterior–anterior) were also obtained to correct for susceptibility-induced distortion in fMRI data. These field maps were collected using a spin-echo echo-planar sequence with the following parameters: flip angle = 90 °, voxel size = 2 × 2 × 2 mm, matrix size = 108 × 108 × 78, Multiband Acceleration Factor = 1, TR = 8330 ms, and TE = 63.40 ms.

T1-weighted (T1w) and T2-weighted (T2w) images were obtained for neuronavigation of the TMS coil during QPS on Days 2 and 3 and for spatial normalization of MRI data. A T1w image was obtained with a magnetization-prepared rapid acquisition with a gradient echo (MPRAGE) sequence with the following parameters: flip angle = 8 °, voxel size = 0.8 × 0.8 × 0.8 mm, matrix size = 224 × 320 × 320, TI = 1000 ms, TR = 2500 ms, and TE = 2.18 ms. A T2w image was obtained with a sampling perfection with application optimized contrast using different flip angle evolution (SPACE) sequence with the following parameters: flip angle = 8 degrees, voxel size = 0.8 × 0.8 × 0.8 mm, matrix size = 224 × 320 × 320, TR = 3200 ms, and TE = 564 ms. Total acquisition time for structural MRI was around 10 min.

On both Days 2 and 3, rsfMRI data were collected to assess the functional changes induced by QPS. These data were acquired with the same acquisition sequence and parameters as the task fMRI data on Day 1. During the scans, participants were required to gaze at a black cross located at the center of a screen for 10 min without any particular thoughts. As with task fMRI on Day 1, a pair of B0 field maps in both phase-encoding directions was obtained with the following parameters: flip angle = 90°, voxel size = 2 × 2 × 2 mm, matrix size = 108 × 108 × 78, Multiband Acceleration Factor = 1, TR = 6240 ms, and TE = 44.00 ms. After each scan, each participant was required to self-evaluate their sleepiness during the scan using the Stanford Sleepiness Scale^35^.

dMRI data were also acquired to assess the microstructural changes induced by QPS. These data were obtained using a 2D single-shot spin-echo echo-planar sequence with the following parameters: voxel size = 2 × 2 × 2 mm, matrix size = 106 × 106 × 75, iPAT reduction factor = 2, Multiband Acceleration Factor = 3, b = 1000 s/mm^2^, number of directions = 30, TR = 5000 ms, and TE = 71 ms. In each session before and after QPS, two sets of dMRI data were collected: one along with the phase-encoding direction anterior–posterior and the other with the reversed phase-encoding direction (posterior–anterior). For each dMRI data, three non-diffusion-weighted images were also obtained in every ten diffusion-weighted images with the same acquisition parameters as the diffusion-weighted images. The total acquisition time for dMRI was approximately 7 min in each session.

### Image Analysis

For the preprocessing of MRI data, unless otherwise noted, the default settings of the preprocessing pipelines of the Human Connectome Project (HCP pipelines 4.3.0, https://github.com/Washington-University/HCPpipelines) were followed, the methods of which are detailed by Glasser et al. (2013)^36^. These pipelines utilize the FMRIB software library (FSL 6.0.5, https://fsl.fmrib.ox.ac.uk/fsl), FreeSurfer (Version 6.0.1, https://surfer.nmr.mgh.harvard.edu/ (Dale et al., 1999; Fischl et al., 1999)), and Connectome Workbench (Version 1.5.0, https://www.humanconnectome.org/software/connectome-workbench) to optimally preprocess MRI data obtained in an HCP-like fashion and perform surface-based analysis (See Supplementary Methods for the detail in preprocessing steps).

After these preprocessing steps, the left M1 responsible for the right FDI from task fMRI data was defined. To define the left M1 responsible for the right index finger, a general linear model (GLM) analysis of task fMRI data was performed with “*TaskfMRIAnalysis.sh*” in the HCP pipelines. This script performs surface-based analysis based on the FEAT tools of the FSL. First, the data were high-pass filtered with a cut-off period of 160 s, and the event timings for tapping the right or left index finger were created with a box-car design in each session, with the contrast tapping the right index finger larger than the left one (RIGHT > LEFT). After applying a 1st-level GLM analysis, a 2nd-level GLM analysis was performed across sessions for each participant. Finally, a Permutation Analysis of Linear Models (PALM) application was used to perform a one-sample t-test towards the contrast RIGHT > LEFT across participants in the MNI standard space.

To quantify the functional changes induced by QPS in both the stimulated and non-stimulated regions, seed-based correlation analyses were conducted towards the rsfMRI data. FC was used for this purpose because this measurement between the stimulated and unstimulated regions was previously shown to be bidirectionally modulated by QPS^18^. The seed was defined as a five mm radius of the peak location identified by the contrast RIGHT > LEFT of the group-level GLM analysis in task fMRI (i.e., the stimulated location) following a previous study^18^. Pearson’s correlation coefficient was calculated between the time series of the seed and that of each vertex or voxel in the whole brain. These coefficients were transformed using Fisher’s z-transformation to conform to normal distribution.

Since a previous study showed that the change in the FC between the left and right M1 was correlated with the magnitude of the after-effect of QPS over left M1^18^, Fisher’s z-transformed Pearson’s correlation was further calculated between the time series of the left and right M1. The right M1 was defined as a five mm radius of the peak location in the contrast LEFT > RIGHT. The functional change of QPS was measured by substituting the FC of these regions post-QPS with that of pre-QPS.

From the dMRI data, the FA and MD values were calculated with a command “*dtifit*” in FSL. The data of both the cerebral cortex and subcortical structures were then warped to the standard space using *NoddiSurfaceMapping* (https://github.com/RIKEN-BCIL/NoddiSurfaceMapping), the method of which is detailed in Fukutomi et al. (2018)^37^). In brief, the data within the cerebral cortex were projected onto the mid-thickness surface and resampled to the MNI standard space with MSMAll (See *Mutlimodal Surface Matching* in Supplementary Methods), whereas those in the subcortical structures were warped to the MNI standard space. All these data were spatially smoothed with a Gaussian kernel of two mm full-width half maximum (FWHM).

To assess changes in the microstructural properties of white matter, the TBSS tool of the FSL was applied^38^. This method creates statistics for group-level voxel-wise analysis in WM, as detailed in our previous study^39^. In brief, the FA map of every individual was nonlinearly registered to the standard space, the main WM structure (skeleton) was created from the spatially normalized FA maps of all participants, and the FA and MD values were projected onto that skeleton.

### Statistical Analysis

The FSL PALM application was used to test the statistical significance of the functional and microstructural changes before and after QPS and the difference in the changes induced by QPS5 and QPS50. Specifically, 10, 000 permutation tests with Threshold-Free Cluster Enhancement^40^ were applied to compare the FC of the left M1 and FA or MD values before and after QPS in each condition and between the changes after QPS5 and QPS50. The Bayes factor was further calculated in each comparison to investigate whether the statistical testing supported the hypothesis that there were no differences in non-significantly different areas. The Bayes factor was calculated using *bayesFactor* (https://github.com/klabhub/bayesFactor). PALM was also utilized to explore the correlation between functional change (i.e., the change in FC between the left M1 and right M1) and the change in FA or MD values. All these p-values were corrected for family-wise errors to control for multiple tests.

To confirm that the neuroimaging results were not affected by other possible confounding factors, such as the stimulation intensity of QPS or the difference in image quality before and after QPS, these measurements were compared with a two-tailed paired t-test for continuous variables and with Wilcoxon signed-rank test for categorical variables. All analyses were performed using the JASP (ver. 0.16.0 for Windows, https://jasp-stats.org/), and *P* < 0.05 was considered statistically significant.

### Data and code availability

The data used in this study will be available by the corresponding authors on reasonable request. All figures and codes are available at BALSA and github, respectively.

## Abbreviations

(QPS): quadri-pulse stimulation
(fMRI): functional MRI
(M1): primary motor cortex
(dMRI): diffusion MRI
(rsfMRI): resting-state fMRI
(FA): fractional anisotropy
(MD): mean diffusivity
(FC): functional connectivity

## Acknowledgements

This work was supported by the Japan Society for the Promotion of Science (Grants-in-Aid for Scientific Research JP18H05523 to K.A. and JP18H01101 to M.J.H., and Grant-in-Aid for Scientific Research on Innovative Areas JP19H05313 to M.J.H.) and Japan Science and Technology Agency (PRESTO JPMJPR17J1 to K.A. and PRESTO JPMJPR19J8 to M.J.H).

## Author contributions

K.I. designed the research; K.I. contributed data; K.I. analyzed the data; K.I. wrote the paper; M.J.H. and K.A provided the feedback on the drafts of the paper; M.J.H. and K.A supervised the work.

