## Supplementary for "Immediate effect of quadri-pulse stimulation on human brain microstructures"

**Supplementary Methods**

Preprocessing

*Structural MRI data*

Structural MRI data obtained on Day 1 (i.e., T1w and T2w images) were preprocessed using "PreFreeSurferPipeline.sh," "FreeSurferPipeline.sh," and "PostFreeSurferPipeline.sh" in the HCP pipelines. Gradient nonlinearity correction was not applied because all participants' heads were placed in the conventional location (i.e., the isocenter) of the MR scanner. First, both T1w and T2w images were transformed to the ACPC space by rigid-body transformation to the Montreal Neurological Institute (MNI) standard space and brain extraction. A T2w image was co-registered to a T1w image, and the bias field was estimated from both the T1w and T2w images. After the bias field correction with this estimated bias field, these images were registered to the MNI standard space, and a modified version of the recon-all command was applied in FreeSurfer to preprocess MRI data with sub-millimeter spatial resolutions. This command generates the boundaries between the cerebrospinal fluid and gray matter (pial surface) and those between the gray and white matter (white surface) and extracts the subcortical structure. The midpoints between the pial and white surfaces were further calculated to create the mid-thickness surface. This surface was utilized to project the fMRI or dMRI data for further surface-based analysis on the cerebral cortex. These surfaces and segmented subcortical volumes were combined and warped to the MNI standard space. Finally,

T1w data were divided by T2w data to derive the myelin map (Glasser and Van Essen, 2011) to be projected onto the mid-thickness surface, which was utilized for multimodal surface matching in this study (See the Multimodal surface matching section below for more details regarding this).

*Functional MRI data*

Both task fMRI and rsfMRI data were first preprocessed using "fMRIVolumeProcessingPipeline.sh" and "fMRISurfaceProcessingPipeline.sh" in the HCP pipelines. First, image distortions caused by the inhomogeneity of the magnetic field were estimated using the corresponding B0 field maps. Then, with these estimated distortions, fMRI data were corrected for EPI image distortions, and motion correction was applied with rigid-body transformation. The corrected fMRI data were co-registered to the T1w image in the ACPC space with boundary-based registration and warped to the MNI standard space. The data in the cerebral cortex were projected onto the mid-thickness surface and resampled to the MNI standard space with spatial smoothing using a Gaussian kernel of 2 mm full-width half maximum (FWHM) along the surface. In contrast, the time series of the subcortical structures were extracted from the subcortical voxels defined from the structural MRI data using FreeSurfer. These voxels were warped to the MNI standard space and spatially smoothed with a Gaussian kernel of 2 mm FWHM.

To further denoise the time series of the rsfMRI data, the "hcp_fix" was applied to the HCP pipelines. This method is based on the FIX application of FSL, which performs spatial independent component analysis towards fMRI data to remove the noise components from these time series[1]. These components were manually re-inspected following the criteria described by Griffanti et al. (2017)[2] and the noise components were regressed outfrom the time series. The mean framewise displacement (FD)[3] was also calculated for each session to quantitatively assess the quality of the rsfMRI data in each session.

*Multimodal surface matching*

To further fine-tune the resampling strategy from the ACPC space to the MNI standard space of the measurements on the mid-thickness, "MSMAllPipeline.sh" was used in the HCP pipelines. This pipeline uses a multimodal surface matching tool[4] of FSL customized for the HCP pipelines (MSMAll[5,6]), which utilizes multimodal features, such as the myelin map and the functional connectivity on each surface, for the registration from the ACPC space to the MNI standard space. After registration, the time-series fMRI data were resampled to the MNI standard space using the resampling strategy defined with this method.

*Diffusion MRI data*

To preprocess the dMRI data, “DiffPreprocPipeline.sh” was applied to the HCP pipelines. First, non-diffusion-weighted images were intensity-normalized and, from these images, image distortions caused by the inhomogeneity of the magnetic field were estimated. With these estimated distortions, the dMRI data were corrected for EPI image distortion and then corrected for eddy current and subject motion. These corrected images were co-registered with the T1w image in ACPC space using BBR. The quantitative measurements of the image qualities in each scan, namely absolute and relative head motions, were calculated using the EDDY QC tools in the FSL.

**Supplementary Results**

<u>Results related to the FC with the left M1 regressing out the effect of mean FD</u>

Analysis of covariance was additionally applied towards the FC of the left M1 obtained with seed-based correlation analysis, setting the mean FD as a covariate of no interest (Supplementary Figure 8). As a result, we observed that after QPS5, the FC of the left M1 was decreased in the right ventral premotor cortex, precentral gyrus, postcentral gyrus, and dorso-medial prefrontal cortex (Supplementary Figure 8A), while after QPS50, it was decreased in the bilateral precentral gyrus, bilateral postcentral gyrus, and bilateral supplementary motor area (Supplementary Figure 8B). The FC of the left M1 increased in the bilateral cerebellum after QPS5 (Supplementary Figure 8C). No significant differences were found in the change in the FC of the left M1 between the QPS5 and QPS50 conditions.

Regarding the FC of the left and right M1 obtained with ROI to ROI analysis, this index significantly decreased after QPS5 ($t = -2.76$, $P = 0.015$), while FC did not significantly change after QPS50 ($t = -1.46$, $P = 0.17$) also after regressing out the effect of mean FD. No significant differences between the QPS5 and QPS50 conditions in the change of FC between these two regions after QPS were observed ($t = -0.75$, $P = 0.47$). We also confirmed that there were no significant correlations between the change in FC after regressing out the effect of mean FD and the change in FA or MD value after both QPS5 and QPS50. These results suggest that the mean FD did not severely affect the results related to the change in the FC of the left M1.

**Supplementary Figures**

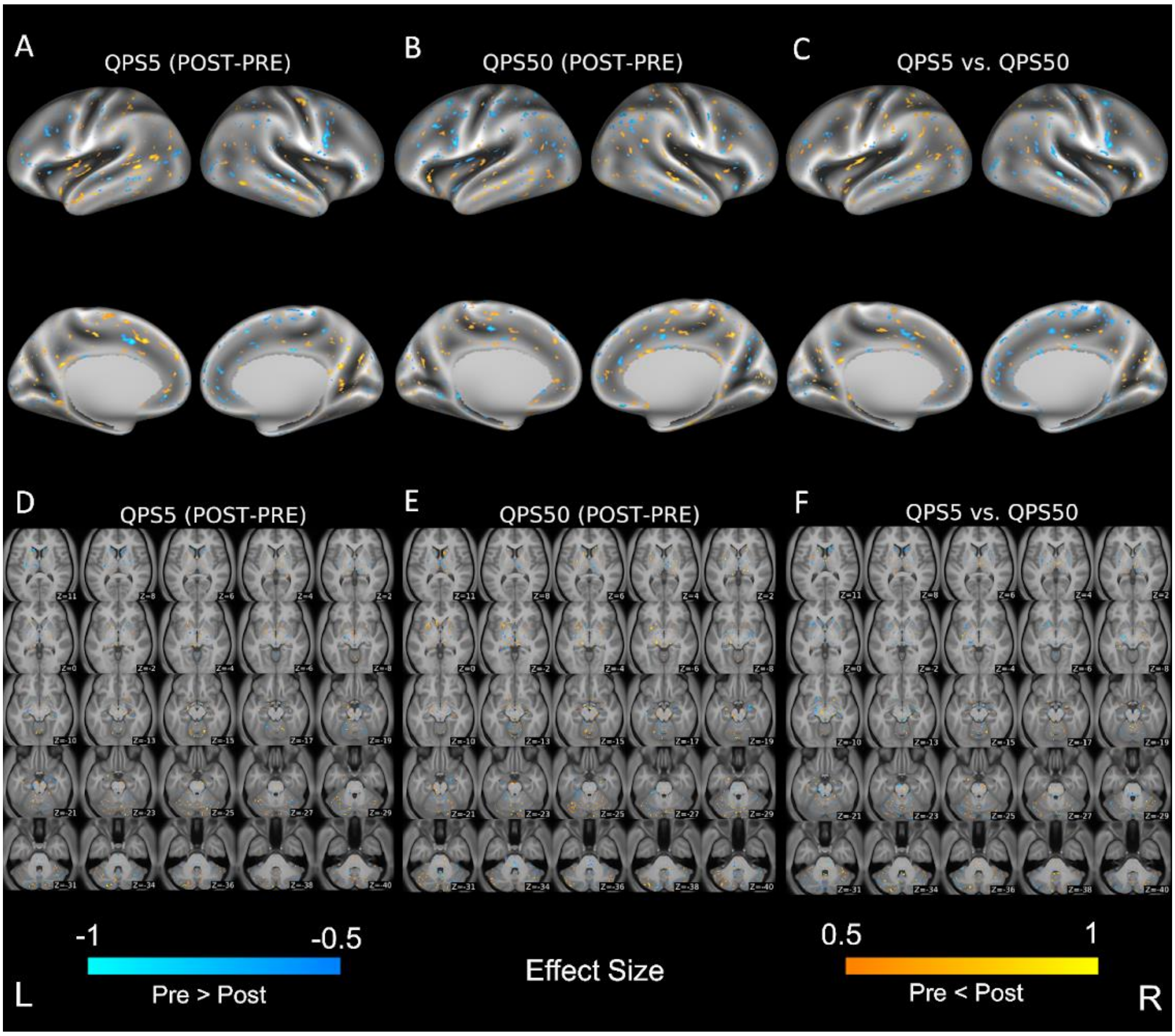


**Supplementary Figure 1.** Differences in FA values of the cerebral cortex and subcortical regions between pre- and post-QPS in QPS5 (A and D) and QPS50 (B and E) conditions, and in changes of FA values across conditions (C and F). The upper panel (A–C) shows the result of surface-based analysis on the cerebral cortex, while the lower panel (D–F) shows that of voxel-based analysis in the subcortical regions. Areas in blue illustrate that the Cohen's d of change in the value after QPS are smaller than -0.5, while those in yellow show that those values are higher than 0.5. Axial slices in (D–F) are shown in accordance with neurological conventions (the left side of the image is of the left of the brain) and are displayed in the MNI coordinates from z = 19 (top left) to z = -57 (bottom right).

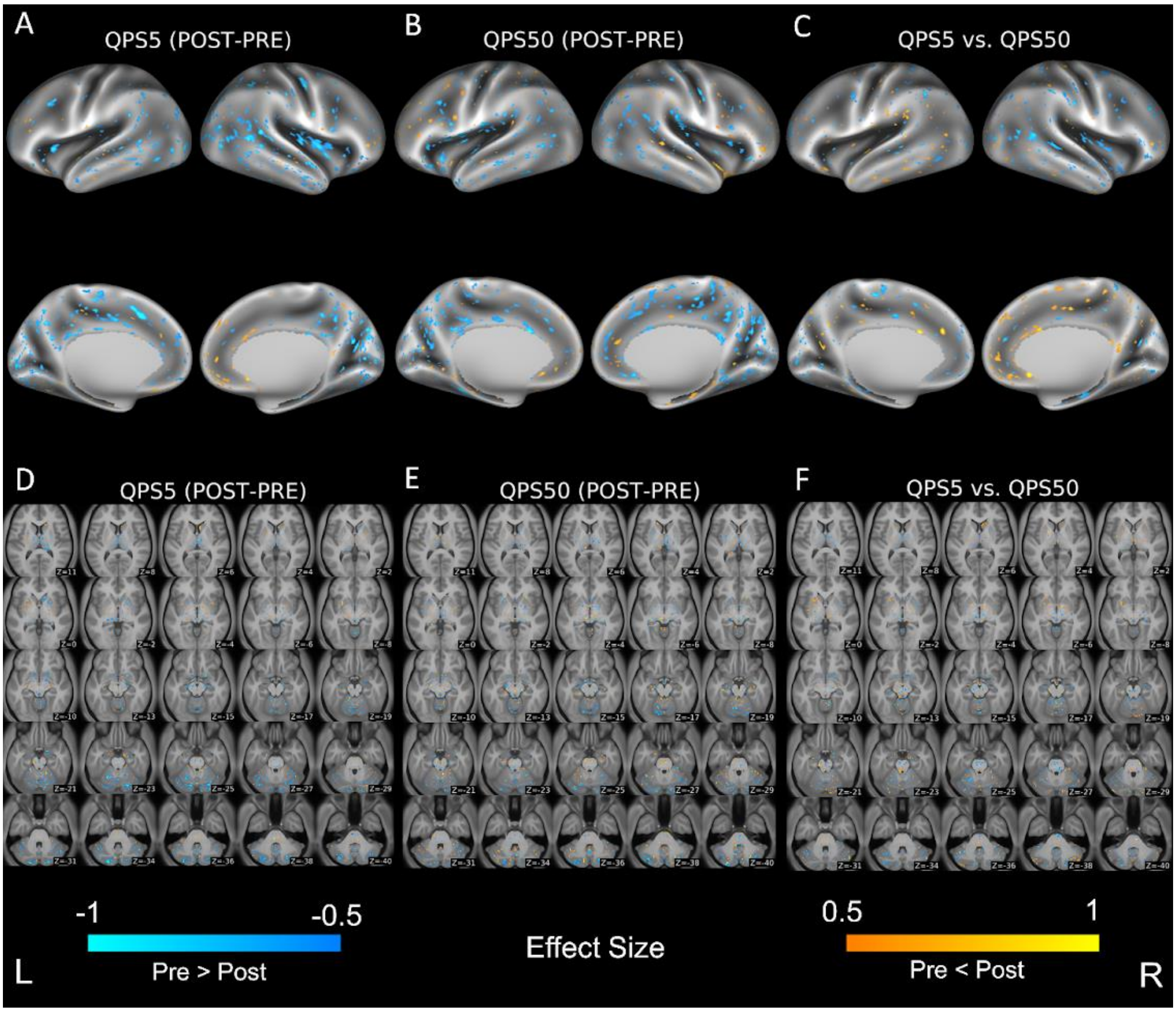


**Supplementary Figure 2.** Differences in MD values of the cerebral cortex and subcortical regions between pre- and post-QPS in QPS5 (A and D) and QPS50 (B and E) conditions, and in changes of MD values across conditions (C and F). The upper panel (A–C) shows the result of surface-based analysis on the cerebral cortex, while the lower panel (D–F) shows that of voxel-based analysis in the subcortical regions. Areas in blue illustrate that the Cohen's d of change in the value after QPS are smaller than -0.5, while those in yellow show that those values are higher than 0.5. Axial slices in (D–F) are shown in accordance with neurological conventions (the left side of the image is of the left of the brain) and are displayed in the MNI coordinates from z = 19 (top left) to z = -57 (bottom right).

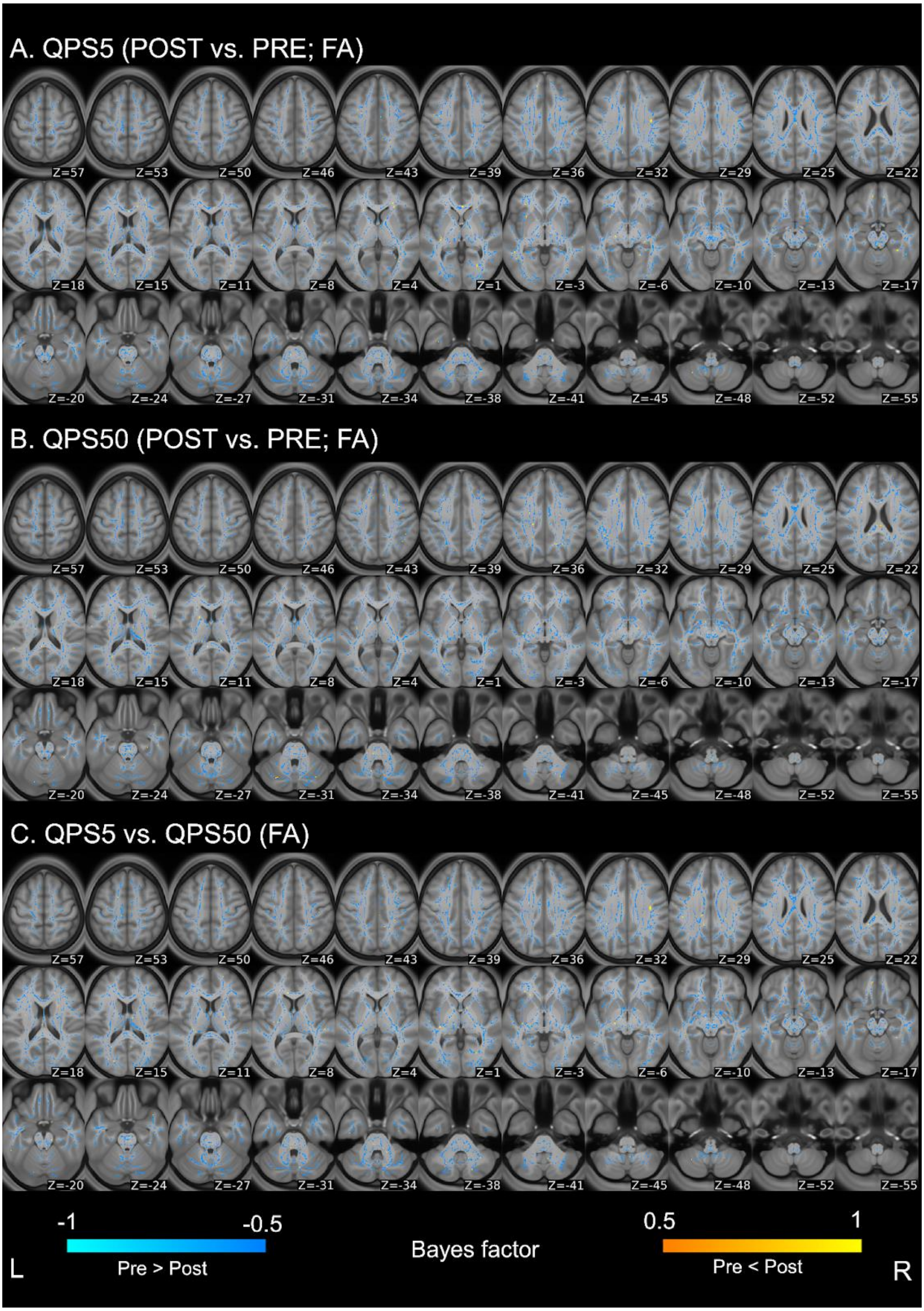


**Supplementary Figure 3.** Differences in FA values of the white matter between pre- and post-QPS in QPS5 (A) and QPS50 (B) conditions, and in the change of FA values across conditions

(C). Areas in blue illustrate that the Bayes factor of change in the value after QPS are smaller than 1/3, while those in yellow show that those are higher than 3. Axial slices are shown in accordance with neurological conventions (the left side of the image is of the left of the brain) and are displayed in the MNI coordinates from z = 57 (top left) to z = -55 (bottom right).

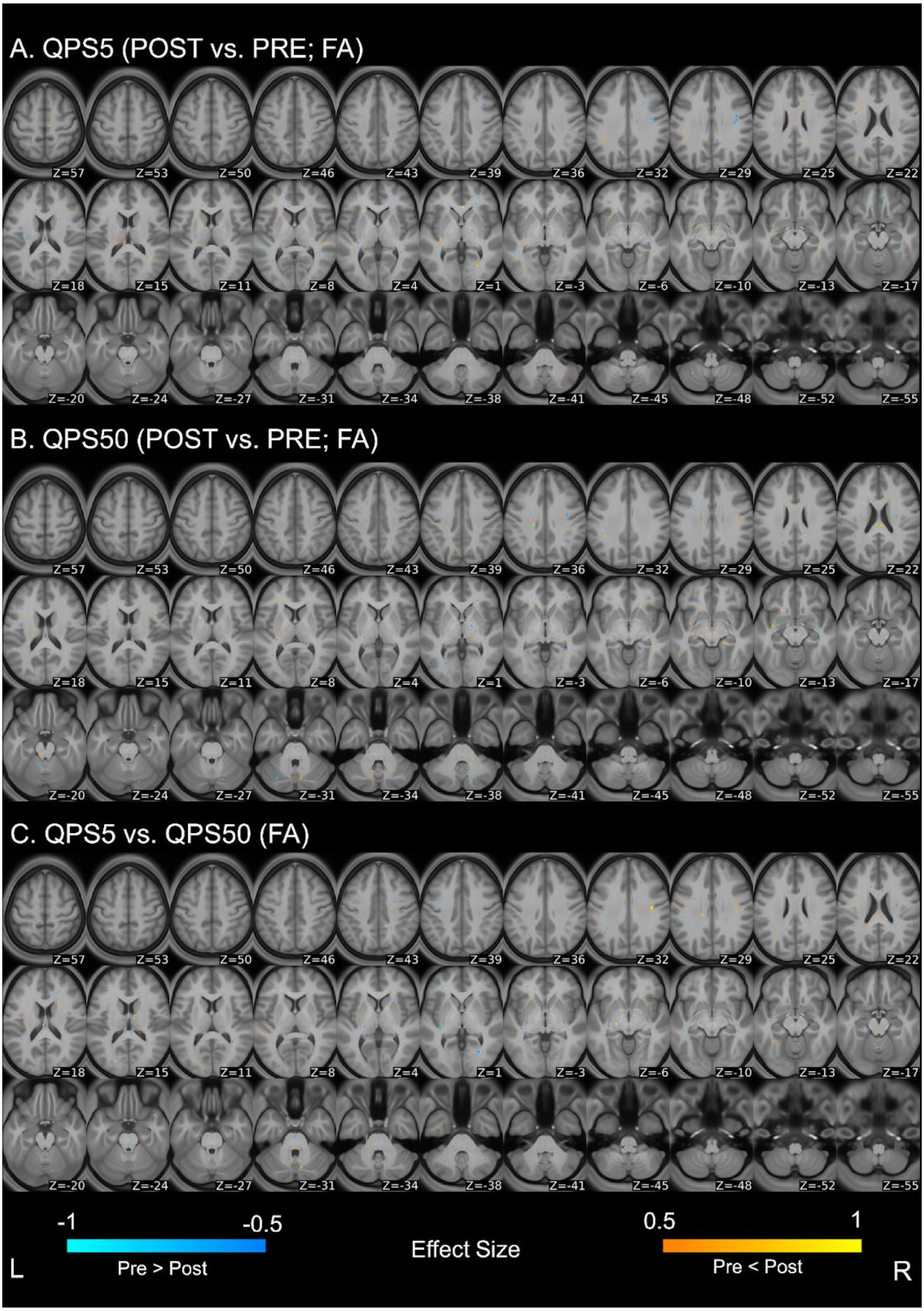


**Supplementary Figure 4.** Differences in FA values of the white matter between pre- and post-QPS in QPS5 (A) and QPS50 (B) conditions, and in the change of FA values across conditions

(C). Areas in blue illustrate that the Cohen's d of change in the value after QPS are smaller than -0.5, while those in yellow show that statistical values are higher than 0.5. Axial slices are shown in accordance with neurological conventions (the left side of the image is on the left of the brain) and are displayed in the MNI coordinates from z = 57 (top left) to z = -55 (bottom right).

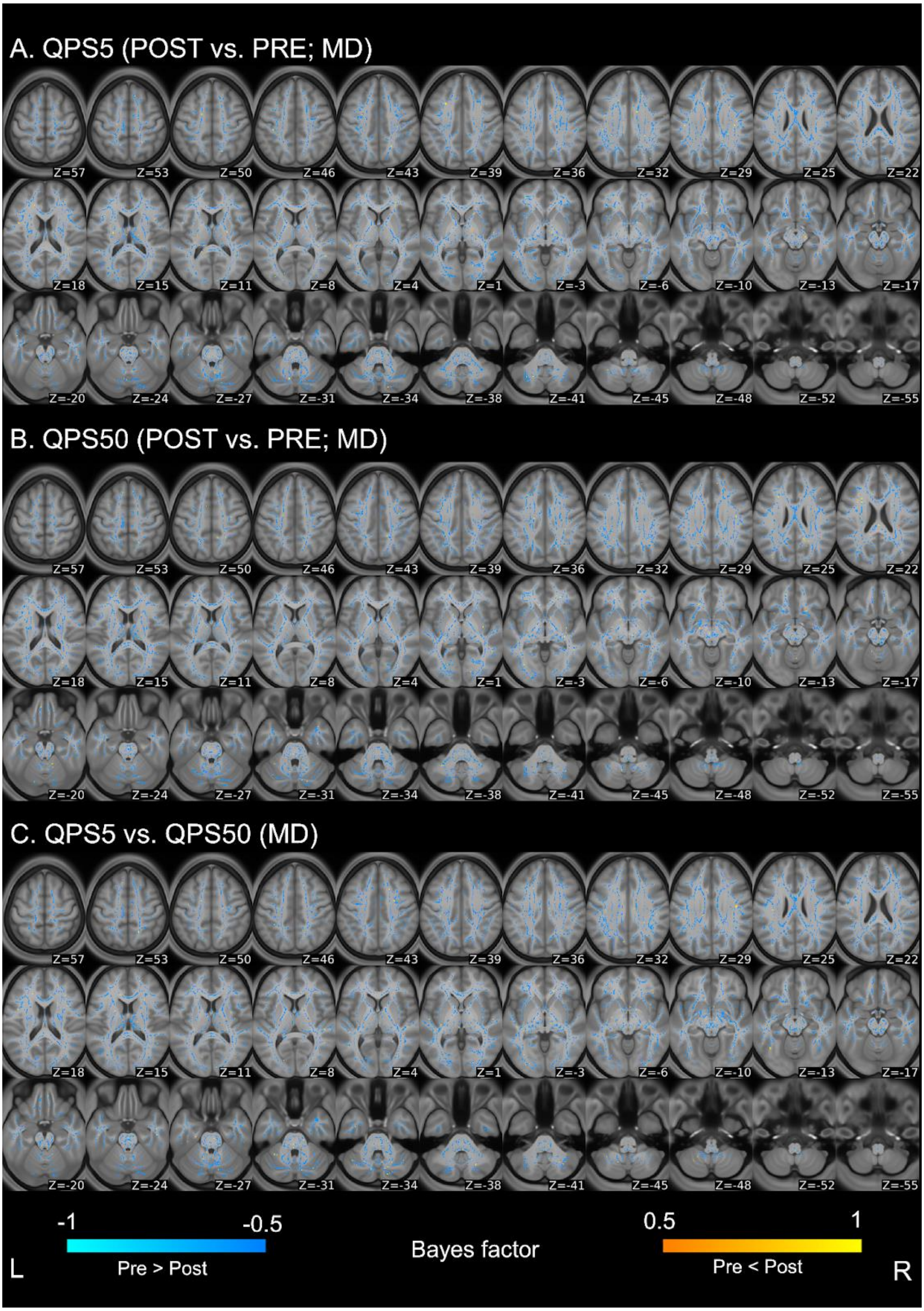


**Supplementary Figure 5.** Differences in MD values of the white matter between pre- and post-QPS in QPS5 (A) and QPS50 (B) conditions, and in the change of MD values across conditions (C). Areas in blue illustrate that the Bayes factor of change in the value after QPS are smaller than 1/3, while those in yellow show that those are higher than 3. Axial slices are shown in accordance with neurological conventions (the left side of the image is of the left of the brain) and are displayed in the MNI coordinates from z = 57 (top left) to z = -55 (bottom right).

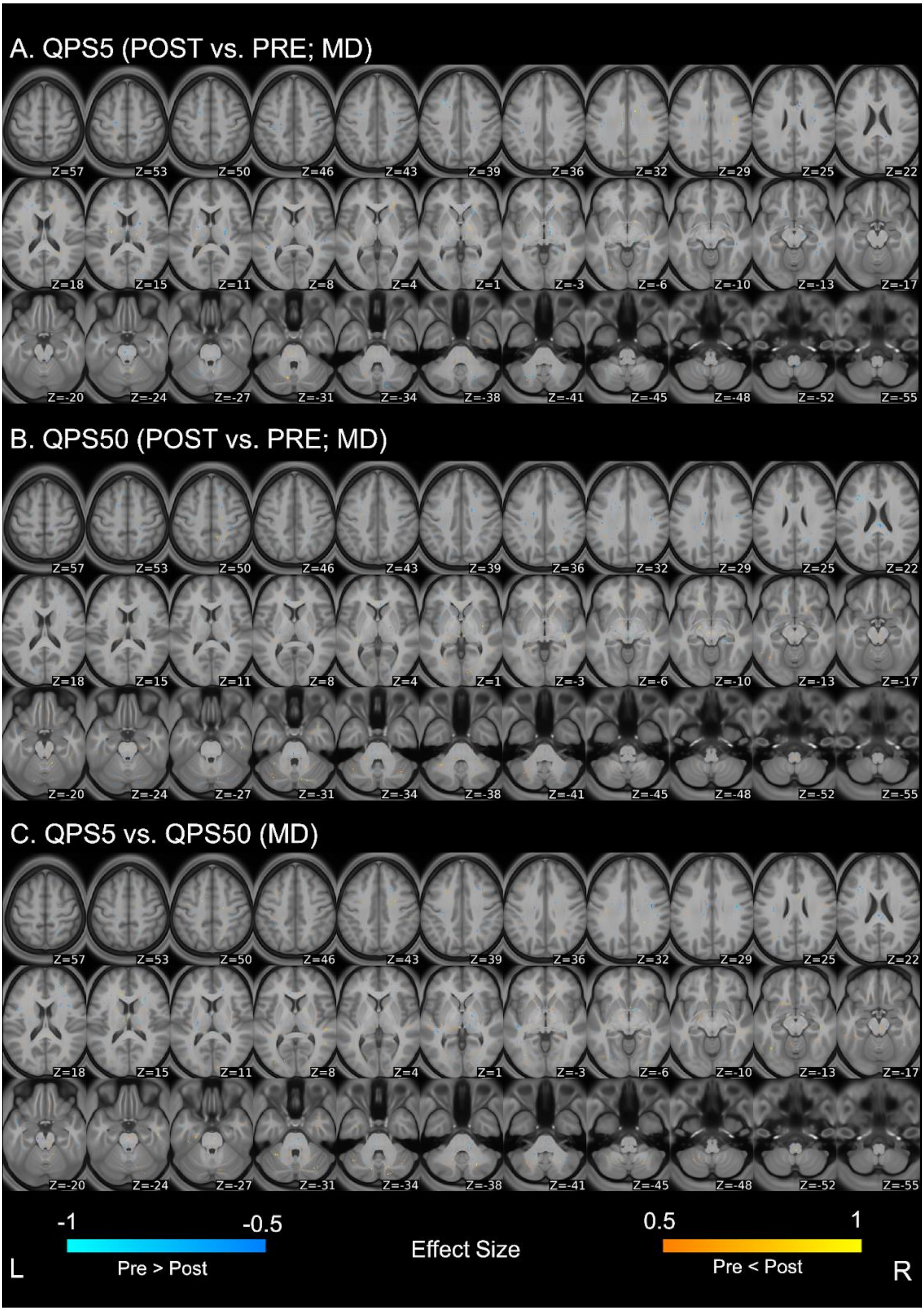


**Supplementary Figure 6.** Differences in MD values of the white matter between pre- and post-QPS in QPS5 (A) and QPS50 (B) conditions, and in the change of MD values across conditions (C). Areas in blue illustrate that the Cohen’s d of change in the value after QPS are smaller than -0.5, while those in yellow show that statistical values are higher than 0.5. Axial slices are shown in accordance with neurological conventions (the left side of the image is on the left of the brain) and are displayed in the MNI coordinates from z = 57 (top left) to z = -55 (bottom right).

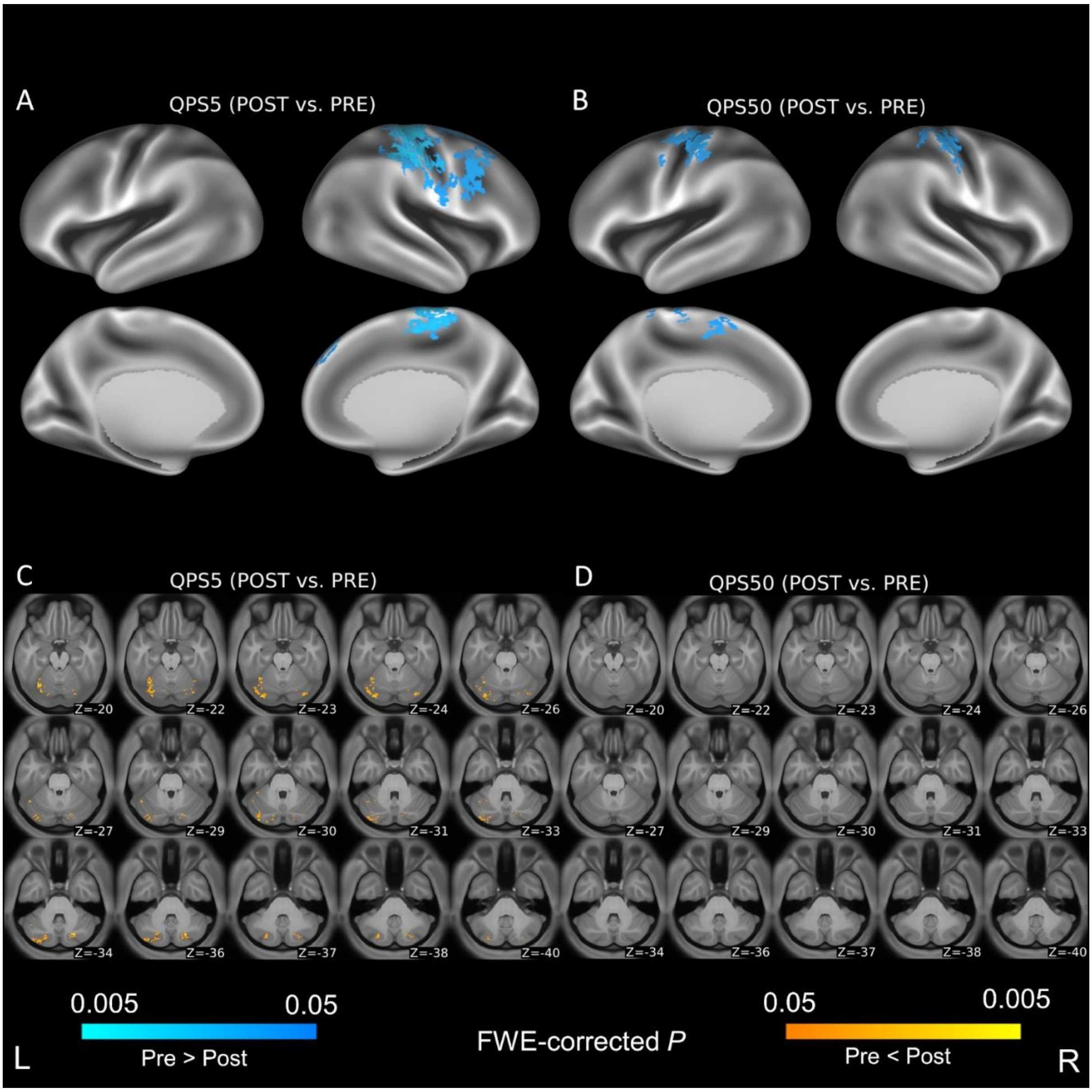


**Supplementary Figure 7.** Differences in the functional connectivity (FC) of the left M1 (i.e, the stimulated region) between pre- and post-QPS in QPS5 (A and C) and QPS50 (B and D) conditions. (A) and (B) show the results of differences with surface-based analysis in the cerebral cortex, while (C) and (D) indicate those with voxel-based analysis in subcortical structures. Areas in blue show that the FC with the left M1 was significantly decreased after QPS, while areas in yellow indicate that that value was significantly increased after QPS. Axial slices are shown in accordance with neurological conventions (the left side of the image is of the left of the brain) and are displayed in the MNI coordinates from z = -20 (top left) to z = -40 (bottom right).

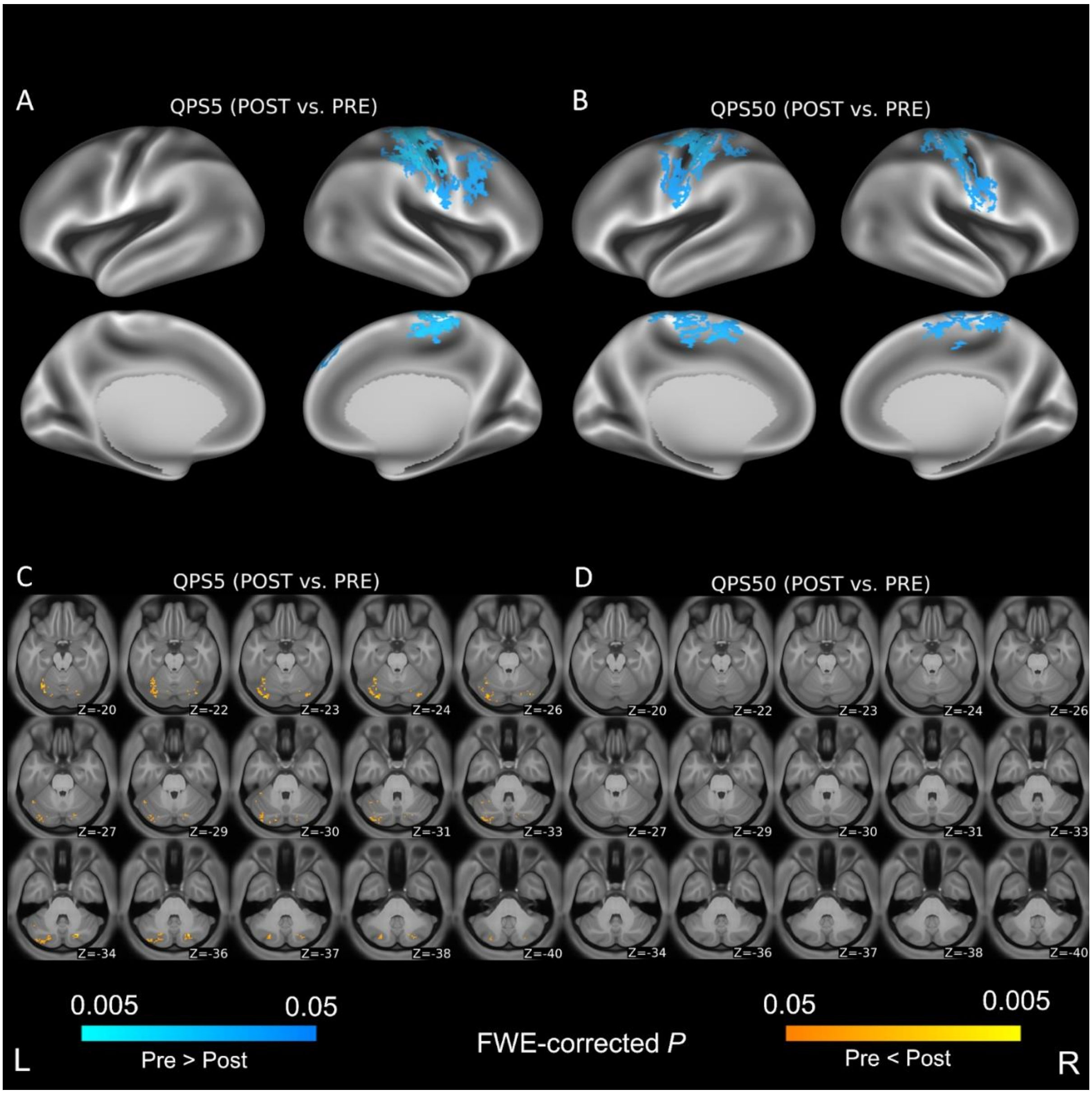


**Supplementary Figure 8.** Differences in the functional connectivity (FC) of the left M1 (i.e, the stimulated region) between pre- and post-QPS in QPS5 (A and C) and QPS50 (B and D) conditions, setting the mean framewise displacement as a covariate of no-interest. (A) and (B) show the results of differences with surface-based analysis in the cerebral cortex, while (C) and (D) indicate those with voxel-based analysis in subcortical structures. Areas in blue show that the FC with the left M1 was significantly decreased after QPS, while areas in yellow indicate that that value was significantly increased after QPS. Axial slices are shown in accordance with neurological conventions (the left side of the image is of the left of the brain) and are displayed in the MNI coordinates from z = -20 (top left) to z = -40 (bottom right).
